# The statistical challenge of finding spontaneous changes in functional connectivity in high-dimensional fMRI data

**DOI:** 10.1101/2020.12.15.422845

**Authors:** Diego Vidaurre

**Affiliations:** Department of Clinical Medicine, Aarhus University; Department of Psychiatry, University of Oxford

## Abstract

An important question in neuroscience is whether or not we can interpret spontaneous variations in the pattern of correlation between brain areas, which we refer to as functional connectivity or FC, as an index of dynamic neuronal communication in fMRI. That is, can we measure time-varying FC reliably? And, if so, can FC reflect information transfer between brain regions at relatively fast-time scales? Answering these questions in practice requires dealing with the statistical challenge of having high-dimensional data and a comparatively lower number of time points or volumes. A common strategy is to use PCA to reduce the dimensionality of the data, and then apply some model, such as the hidden Markov model (HMM) or a Bayesian mixture of distributions, to find a set of distinct FC patterns or states. The distinct spatial properties of these FC states together with the time-resolved switching between them offer a flexible description of time-varying FC. In this work, we show that in this context PCA can suffer from systematic biases and loss of sensitivity for the purposes of finding time-varying FC. To get around these issues, we propose a novel variety of the HMM, the HMM-PCA, where the states are themselves PCA decompositions. Since PCA is based on the data covariance, the state-specific PCA decompositions reflect distinct patterns of FC. We show, theoretically and empirically, that fusing dimensionality reduction and time-varying FC estimation in one single step can avoid these problems and outperform alternative approaches, eventually facilitating the quantification of transient communication in the brain.

**Author summary:** We show that PCA, although widely used in practice, can introduce important biases and loss of sensitivity in the estimation of time-varying functional connectivity on high-dimensional fMRI data. We discuss these limitations and propose a new method that, by performing dimensionality reduction and time-varying functional connectivity estimation in one single step, can effectively overcome these limitations.

## 1 Introduction

When we image the brain of passive subjects with fMRI, the measured signals exhibit strong correlations even between areas that are far apart in the brain [1, 2]. These patterns of *resting-state* correlation, referred to as functional connectivity (FC), are interpreted as a sign that these regions are somehow engaged together in relation to one or more brain processes [3]. It is now widely recognised that FC holds important relations to mental and clinical phenotypes, is reliably subject-specific (i.e. reproducible across scanning sessions), and is also hereditary [4, 5, 6]. However, the mere existence and interpretability of within-session modulations in FC is, at least in fMRI, more controversial [7, 8]. An important reason for this dispute is the scarcity of time samples and the very high dimensionality of the data. In this context, quantifying modulations of FC within session is a challenging statistical problem because these changes (if they exist) are by definition spontaneous and have no obvious behavioural reference [9].

One possible strategy to quantify time-varying FC is the use of sliding-windows, where a correlation matrix is computed for each window in the data [10]. Although attractive because of its simplicity, this method suffers from an important problem of statistical variability in the estimation, and there is no obvious way to disentangle actual changes in the data from fluctuations caused by statistical noise [11]. For this reason, methods that boost the statistical power by using the entire data set in the estimation are sometimes preferred. One such method is the Hidden Markov model (HMM), which assumes that the data can be reasonably modelled using a discrete number of FC states with Markovian dynamics. In this case, the Markovian property means that the model accounts for the state dynamics using a probability matrix that encodes the probability of transitioning between each pair of states – but without considering the previous history of state activations [12, 13]. A straightforward variant of the HMM is to let each state be modelled as a Gaussian distribution with the mean pinned to zero in order to prioritise changes in covariance [14, 8]. A simpler alternative is a Bayesian mixture of distributions [13], which has no transition probability matrix and therefore ignores the temporal structure of the data.

Unfortunately, neither the HMM nor the Bayesian mixture of Gaussian distributions are easily applicable when the data dimensionality (the number of voxels) is too high in relation with the number of time points (volumes). The two most common approaches for reducing dimensionality in this context are using an anatomical parcellation [15] and applying independent component analysis (ICA) [3]. These produce a number *n* of regions or components (typically a hundred or more), so that an FC matrix has *n*(*n* – 1)/2 different parameters. Often, this is still high enough for the HMM or Bayesian mixture inference to overfit and produce degenerate solutions in many data sets. For this reason, PCA is usually carried out on the ICA-derived or parcellated time series so that the HMM (or alternative method) is run on an even lower-dimensional space [16].

Whereas this two-step method works reasonably well in practice for many (but not all) fMRI data sets, having PCA and the HMM estimation as two separate steps is suboptimal. This is because the PCA step specialises in maximising explained variance, and is not designed for the final goal of the analysis. Furthermore, the use of PCA can unknowingly introduce important biases on the estimation. In this paper, we discuss these issues in detail and propose an alternative new model that bypasses these problems: an HMM where each state is a PCA decomposition. We refer this model to as *HMM-PCA*. Critically, because the computation of PCA is based on the data covariance [17], the state-specific PCA decompositions reflect distinct patterns of FC, effectively fusing dimensionality reduction and time-varying FC estimation in one single step. Figure 1 presents a graphical illustration of both the HMM with Gaussian states over principal components (HMM-Gaussian, top), and the HMM-PCA (bottom).

**Figure 1:**
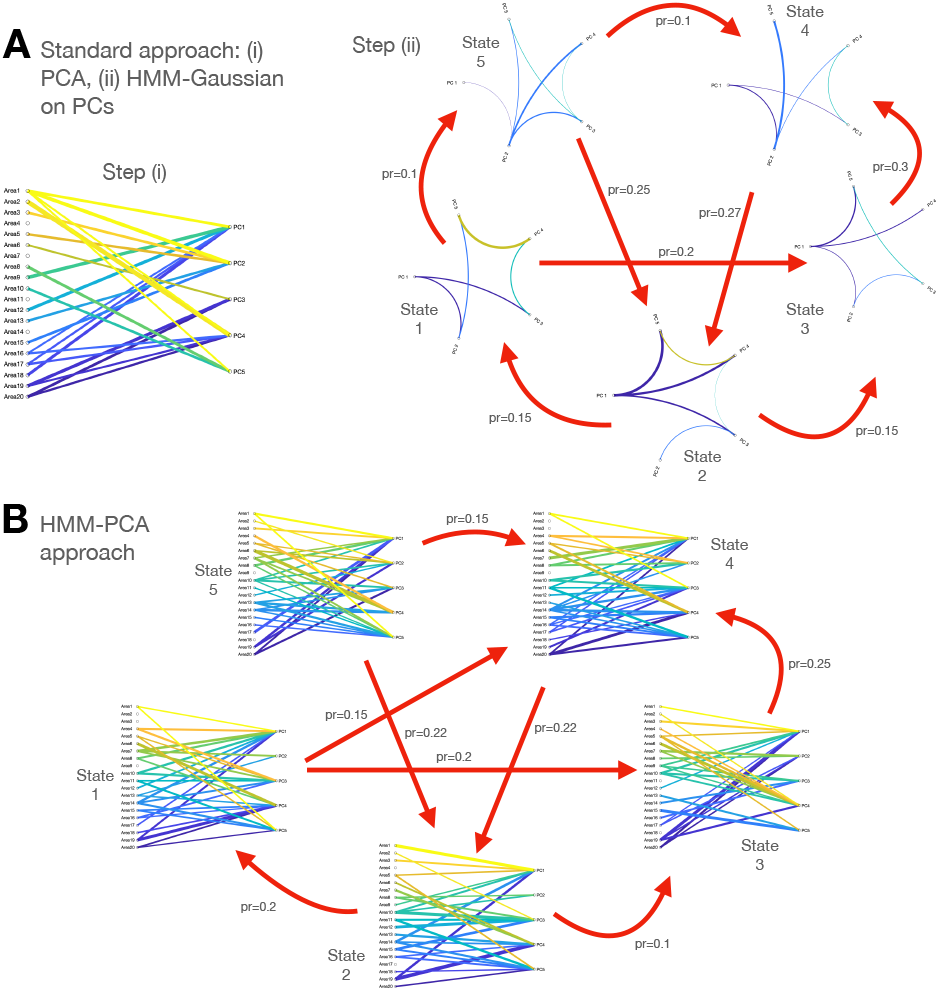
Two different approaches for the estimation of time-varying FC on high-dimensional fMRI data: **A**. PCA is first used as a dimensionality reduction step, blindly to the purpose of estimating timevarying FC; then, some state-based model (like the hidden Markov model) is run on the first principal components (PC). **B**. The HMM-PCA approach, where each state is a different PCA decomposition, is run directly on the high-dimensional data; given that the computation of PCA is based on the data covariance, different PCA decompositions capture different patterns of FC.

## 2 Materials and Methods

### 2.1 The problem of estimating time-varying FC in high dimensions

Let 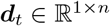 be the multivariate source signal at volume (time point) *t* = 1…*T*, so that ***D*** denotes the data concatenated for all sessions and subjects – although what follows can be also applied at the single-subject level provided that we have a sufficient number of volumes. Here, *n* corresponds to anatomical parcels or ICA components, referred to generically as regions. A standard estimation of functional connectivity (FC) for subject *j* is an *n* × *n* matrix **C** containing the Pearson’s correlation coefficient for each pair of regions. Formally, the question at hand can be posed as: can we find differences between matrices **C**(*t*_1_) and **C**(*t*_2_), defined as *instantaneous* FC matrices at time points *t*_1_ and *t*_2_, for at least one pair of time points *t*_1_ and *t*_2_ belonging to the same scanning session?

One way to approach this problem is the use of the Hidden Markov Model (HMM), which describes the data in terms of a finite number *K* of states that activate or deactivate throughout the scanning time. The state time courses, reflecting these activations, are in the form of probabilities *P* (*x_tk_* = 1|***D***) = *γ_tk_*, where *x_tk_* = 1 means that the state *k* is active at time point *t*, and *γ_tk_* is then the estimated posterior probability of *x_tk_* = 1 given the data. The HMM is a generic model where the states can be described using any family of probability distributions. Within this framework, we can define the states here as covariance matrices **Σ**^(*k*)^; i.e. as Wishart distributions or, equivalently, as zero-mean Gaussian distributions [8]. We refer to this approach as *HMM-Gaussian.* Alternatively, the Bayesian mixture of Gaussian distributions dispenses with the transition probability matrix, thus ignoring the temporal structure of the data and treating the time points (volumes) as independently distributed and exchangeable [13]. We shall refer to this model as *Mix-Gaussian.*

Because *n* is typically large in comparison to *T*, PCA is typically used as an intermediate dimensionality reduction step. This way, HMM-Gaussian (or Mix-Gaussian) is estimated on ***y**_t_* = ***d**_t_**W***, where 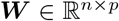 represents a PCA decomposition and *p* is the number of principal components (PCs). The estimated FC matrices are then low-dimensional, 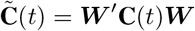. Note that, across the entire PCA-reduced data set, 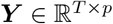, the *p* PCs are by construction orthogonal, but there may exist transient departures from orthogonality. These transient departures from orthogonality, encoded by the HMM parameters *γ_tk_* and **Σ**^(*k*)^, can be regarded as changes in FC – insofar, of course, as the state transitions are not solely caused by relative modulations in variance between regions.

### 2.2 HMM-PCA: a new model for estimating time-varying FC

We now introduce the HMM-PCA mathematically. Various of the elements this model are analogous to the (Bayesian) mixture of PCA analysers introduced by Tipping and Bishop [18] (here referred to as *Mix-PCA*), which, similarly to Mix-Gaussian, does not account for the temporal structure of the fMRI data. In brief, the main read-outs of HMM-PCA and Mix-PCA are (i) a set of *K* states, each characterised by a PCA decomposition; and (ii) the corresponding state time courses *γ_tk_*, which encode the probability of each state *k* to be active at each time point *t*. In the case of HMM-PCA, a matrix with transition probabilities between states is also estimated.

Both Mix-PCA and the HMM-PCA are based on Bayesian PCA [19, 20], which formulates classic PCA within a probabilistic framework. Bayesian PCA assumes the following distribution:

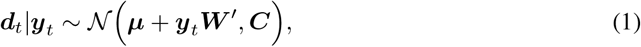

where the covariance is given by ***C*** = ***WW′*** + *σ*^2^***I***, and ***I*** is the identity matrix. The density function is therefore

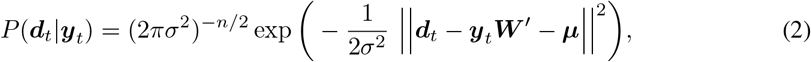

where ***y**_t_* is assumed to be Gaussian distributed, and ║·║ is the Euclidean norm. Since we are not interested in modelling transient changes in amplitude but just in FC, we here modify this model, as opposed to [18], so that ***μ*** = **0**: that is, we will not model changes in amplitude explicitly.

To model the data, the Mix-PCA model uses *K* different PCA projections, 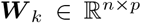, and their corresponding noise variance estimations 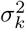. The state covariance matrices are denoted as 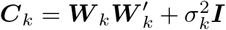. The prior probability for each state to have generated unseen data is given by *π_k_*, which also needs to be estimated. Therefore a second set of latent variables ***x**_k_* = (*x*_1*k*_,…, *x_Tk_*) is required, such that *x_tk_* = 1 if the *k*-th component (or state) is responsible for having generated the observed data at time point *t*, and *x_tk_* = 0 otherwise. We refer to ***X*** = [***x***_1_… ***x**_K_*], as the state time courses. The posterior probabilities *P*(*x_tk_* = 1|***d**_t_, π_k_*) = *γ_tk_* can be estimated, for example, using the EM algorithm [18, 20]. Note that the estimation of *γ_tk_* only depends on the data at time point *t* and on *π_k_*, ignoring the data temporal structure.

In the HMM-PCA case, we instead have the state latent variables ***x**_k_* modelled as an order-1 Markovian process, with prior probabilities *P*(*x*_*tk*_1__ = 1|*x*_(*t*−1)*k*_2__ = 1) = Θ_*k*_2_*k*_1__. Here, **Θ** is the transition probability matrix, which models the average probabilities of transitioning from one state to another – and must be estimated as well. This way, the posterior probabilities (namely, the state time courses) are defined as *P*(*x_tk_* = 1|*x*_(*t*−1)_, ***d**_t_*, **Θ**) = *γ_tk_*. These are estimated using the forward-backward equations, given the likelihood for each HMM state (here, PCA decomposition) at *t* [12].

In order to use the EM algorithm to solve the HMM-PCA problem, following [18] we formulate the (expected) complete log-likelihood for each *k* and time point *t*:

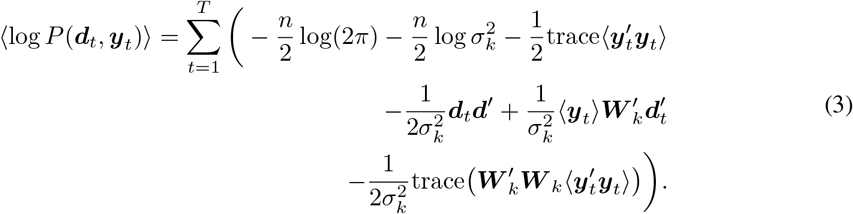

where 〈·〉 denotes expectation, and

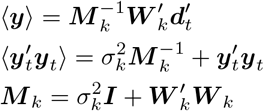

Similarly to the Mix-PCA model, the EM updates for ***W**_k_* and 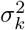 are coupled. Using the same estimation of the intermediate variable ***M**_k_* for both, the new parameter estimates for these variables become:

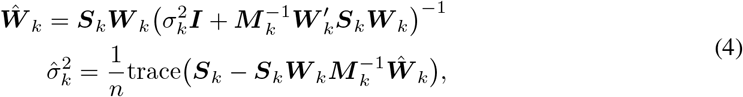

where

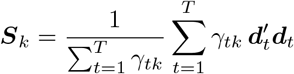

is the state-specific (sample) covariance matrix in the original space, such that the aggregated sample covariance matrix 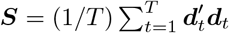 can be expressed as a weighted average of ***S**_k_*. For the full EM algorithm, it only remains the estimation of the state activation probabilities *γ_tk_*, the transition probability matrix **Θ**, and the initial probabilities ***π*** for the first time point of the scans. The forwardbackward equations for estimating *γ_tk_* as well as the update rules for **Θ** and ***π*** are equivalent to any other HMM given the expected log-likelihood in 4, and can be found elsewhere [13, 21].

### 2.3 Limitations of the two-step approaches

Previous work has shown that the HMM, when ran on PCA time series, can produce useful representations of the data [16]. However, this approach suffers from two limitations: (i) a bias towards the lower-order PCs (that is, those explaining less variance), and (ii) a loss of statistical efficiency in detecting time-varying FC when there is time-varying-FC-relevant information in the discarded PCs. These limitations, which we discuss next, equally apply to other probabilistic models and clustering methods such as Mix-Gaussian when applied on PCA time series.

#### 2.3.1 Bias towards low-order PCA components

The application of PCA systematically alters the HMM or Bayesian mixture inference, biasing it towards low-order PCs and introducing a factor of arbitrariness in the inference. This issue occurs even when we keep all PCs and retain 100% of the variance.

When states are described as Gaussian distributions, the HMM (or Bayesian mixture) inference is scale-invariant. That is, as far as we use the same random seed in the initialisation of the inference, the estimation of the state time courses *γ* will not be affected if we multiply the time series of any given region by any random scalar. Mathematically, this can be expressed as

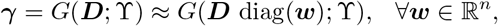

where *G*(***D***; **ϒ**) represents the HMM inference process given some specification of hyperparameters **ϒ** (e.g. the number of states). That is, the inference remains unaltered after rescaling the regions’ time series by any vector ***w***, regardless of the specific values of such vector. Intuitively, the reason is because the state-specific covariance matrices **Σ**^(*k*)^ can adjust their diagonal (i.e. their variance) to compensate for this global scaling with no effects in the inference. (This is as far as the prior distribution of the covariance matrices acknowledges this scaling; if not, there might be small changes in the inference but rarely substantial provided that we have enough data).

However, the HMM (or Bayesian mixture) inference is not rotation-invariant, and, in particular, it is not invariant to a PCA rotation:

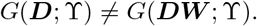

When we apply PCA on the data, ***Y*** = ***DW***, the columns of ***Y*** are ordered according to its variance, such that the first column of ***Y*** (the first eigenvector) has the highest variance and the last column has the lowest variance. Because of the scale-invariance property of the HMM inference, however, these variances will be ignored. Intuitively, this means that the low-order PCs (which explain less variance in the original data) are given in principle the same weight in the inference than the high-order PCs (which explain more variance in the original data). In practice, that results in a distortion of the estimates with respect to the original data, which will become more drastic as we include more and more low-order PCs.

For example, let us consider one given subject from the Human Connectome Project (HCP) data set [22], eight randomly-chosen brain parcels from the (100 regions) Schaefer parcellation [23], and a fixed initialisation of the algorithm (i.e. initialising the inference with exactly the same starting state time courses, so there is no randomness anywhere throughout the inference). Figure 2A shows the estimated state time courses for the original data (top), the state time courses estimated after scaling each channel randomly (middle), and the state time courses obtained from a PCA decomposition where we kept all components so that there is no loss of information (bottom). As observed, PCA-rotating the data changes the estimation, whereas scaling does not.

**Figure 2:**
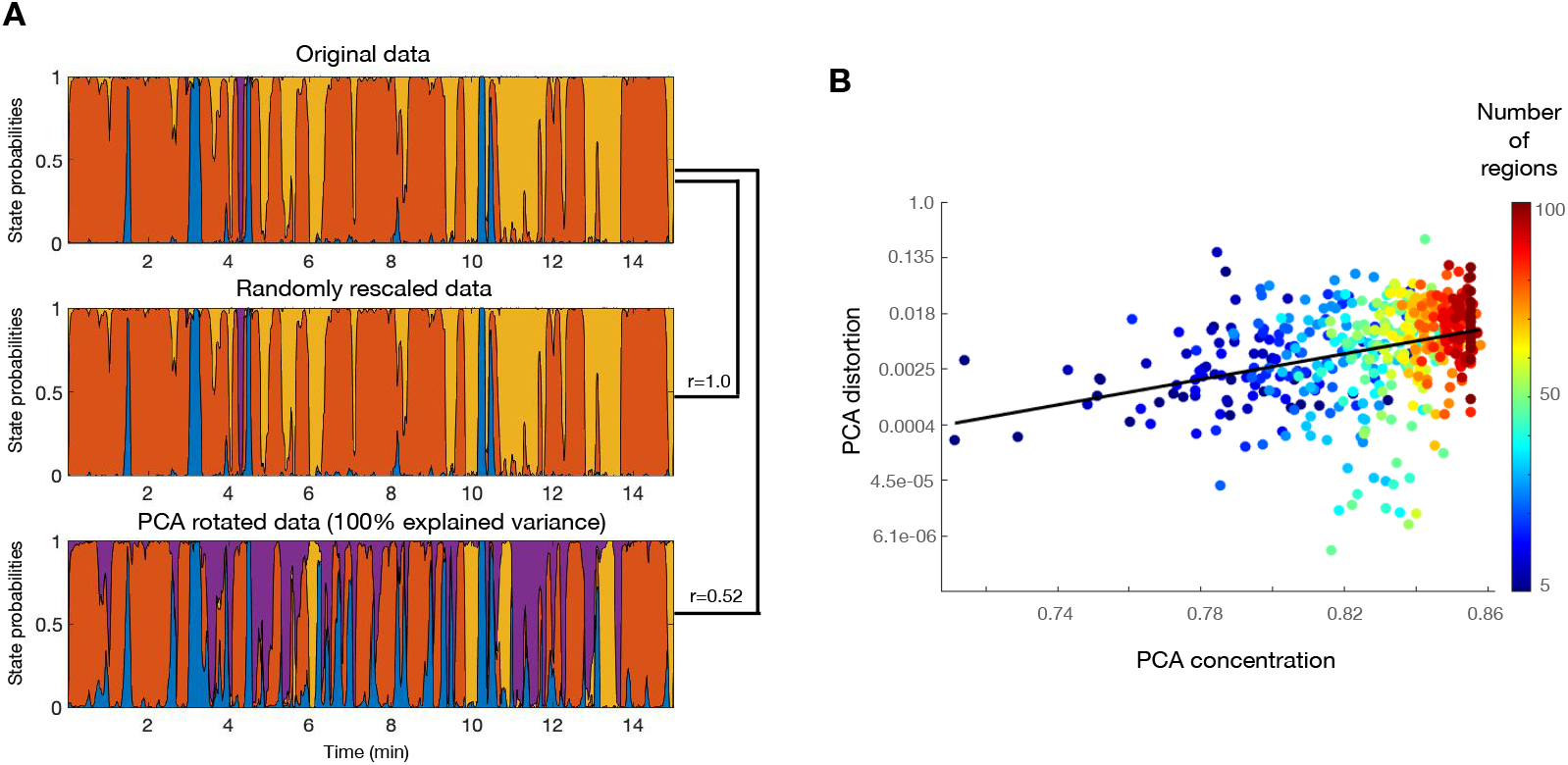
PCA introduces a bias on the HMM (or Bayesian mixture model). **A**. The HMM inference is scale-invariant, but not PCA-rotation-invariant. State time courses produced by the HMM inference for one HCP subject on the original parcellation space (top), after applying a random scaling of the data (middle), and after PCA rotation with no loss of variance (bottom). Each colour represents a different state, so that the coloured areas indicate the probability of activation for the states across the session. The similarity between the different runs, expressed as Pearson’s correlation coefficients, are expressed on the right. **B**. The extent of this bias (PCA distortion) is logarithmically related to how concentrated is the variance on the first PCs (PCA concentration); that is, the more correlated are the regions on the original data, the stronger will be the bias introduced by PCA.

Furthermore, this bias is related with the number of regions and how concentrated is the variance on the first PCs. This is shown in Figure 2B. The amount of *PCA distortion* (Y-axis) is here quantified as one minus the correlation between the state time courses obtained on the original data vs. those obtained on the PCA projection. A measure of PCA *concentration* (X-axis) is given by the average cumulative explained variance across PCs; for example, if the areas were perfectly correlated then the first PC would explain all the variance (1.0) and the cumulative explained variance of all PC would be 1.0, in which case the average −the PCA concentration– would be exactly 1.0; on the opposite case, if the regions were orthogonal (uncorrelated), then the cumulative explained variance of the *j*-th PC is *j/n*, and the average is exactly 0.5. In each run, we sampled a number (between 5 and 100) of regions from the Schaefer parcellation and run the HMM on both the original and the PCA-projected data (with no loss of variance). Figure 2B shows that there is a logarithmic relation between PCA concentration and PCA distortion across HMM runs, suggesting that the more correlated the regions are, the stronger is the bias introduced by PCA.

In summary, although PCA is an acceptable approximation in practice, it can also arbitrarily distort the time-varying FC estimates towards the lower-order PCs. This effect will be more pronounced when the regions are more correlated – i.e. when the proportion of variance explained by the different PCA components is less equally balanced.

#### 2.3.2 Loss of sensitivity

The previous section discussed the biases of estimating HMMs and Bayesian mixtures of Gaussian distributions on PCA-reduced data. We now discuss the loss of sensitivity in detecting time-varying FC when the discarded PCA components contain time-varying-FC-related information. Given that within-session modulations in FC are bound to be subtle [8, 7], it is quite possible that such modulations will indeed occur in lower-order PCs. Since this is not straightforward to show without access to the ground truth, we used simulations where, as it happens in fMRI, the temporal modulations of covariance are not very large. These simulations demonstrate empirically that HMM-PCA can outperforms competing models if there is time-varying FC in lower-order PCs. Secondarily, these simulations stress the importance of acknowledging the temporal nature of the data, which is ignored by Mix-PCA.

We generated data from two different simulation schemes. In both cases, the data were generated from a low-dimensional space and projected to the dimension of the observed data (*n* = 10), where some observational 10-dimensional white noise was added. (That is, the data is low-rank up to the observational noise). The nature of this projection varies according to two different states, which transitions are organised as an order-1 Markovian process with transition probabilities, *P*(*x*_*tk*_1__ = 1|*x*_(*t*−1)*k*_2__) = .9615 if *k*_1_ = *k*_2_ and *P*(*x*_*tk*_1__ = 1|*x*_(*t*−1)*k*_2__) = .0385 otherwise; that is, the probability of remaining in the same state is 25 times higher than that of transitioning. We sampled 10 sessions of 1000 data points each, which is a common data set size; for example, this would correspond to a data set with sampling rate or TR≈1.1s, and 15min worth of data per session. We generated data from a Markovian process for simplicity, but the fact that the generative process aligns with the Markovian assumption of the HMM does not result in any loss of generalisation for the main point of the simulations.

In Scenario 1, we separately sampled each of the latent dimensions from a zero-mean Gaussian distribution, so that these are approximately orthogonal. We denote the dimension of the latent space as *p*_0_, and the generated latent data as ***Y***_0_. Two different separate cases are considered: in the first, we have *p*_0_ = 2, where we sampled two latent variables with standard deviations 2.0 and 1.5; in the second, we have *p*_0_ = 3, with respective standard deviations of 2.0, 1.5 and 1.0. We then sampled three random (standard Gaussian-distributed) projection matrices *A, A*_1_, 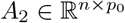, and generated the observed data as ***S*** = ***Y***_0_ (*A* + *A*_1_) + *ϵ* when state 1 is active, and ***S*** = ***Y***_0_ (*A* + *A*_2_) + *ϵ* when state 2 is active. Observational noise e is set to have zero mean and standard deviation equal to 0.001.

In Scenario 2, we based the sampling on actual fMRI data from the HCP (see above). In particular, we randomly chose *n* =10 regions from the data and computed the data covariance matrix ***C***, which we then eigendecomposed into its principal components: ***C*** = ***V′EV***, where ***E*** is a diagonal matrix of eigenvalues and ***V*** contains the corresponding eigenvectors. We used these to create two covariance matrices ***C***_1_ and ***C***_2_, which were then used to sample the data for each of the two states given a multivariate Gaussian distribution with zero mean. As with the other scenario, we considered two cases: *p*_0_ = 2 and *p*_0_ = 3. In either case, the first eigenvector of both ***C***_1_ and ***C***_2_ was set to be the first eigenvector of ***C***, thus corresponding to the time-invariant component of FC. In the *p*_0_ = 2 case, the second eigenvector of ***C***_1_ and ***C***_2_ were assigned each a different permutation of the second eigenvector of ***C***, and the rest of eigenvectors were set to zero. In the *p*_0_ = 3 case, the second and third eigenvectors of ***C***_1_ and ***C***_2_ were assigned different permutations of the second and third eigenvector of ***C***, and the rest of eigenvectors were set to zero. These latter eigenvectors, therefore, represent the time-varying components of the FC. Again, additive observational noise *ϵ* was set to have zero mean and standard deviation equal to 0.001.

We repeated the simulations 50 times, and estimated HMM-PCA, Mix-PCA and HMM-Gaussian models, which were all set to use *p* = 2 components, and, for simplicity, the right number of states *K* = 2. Therefore, there is loss of information only in the *p*_0_ = 3 cases. For illustration, Figure 3A shows one specific instance for Simulation 1 and *p* = *p*_0_ = 2, where the HMM-Gaussian approach misses some of the swiftest state changes, and Mix-PCA, on the other hand, appears to be noisier due to the lack of consideration for the temporal structure of the data. Figure 3B shows the complete results. Here, accuracy corresponds to the proportion of time where the corrected state was guessed. Each dot corresponds to one instance of the simulations, representing how the HMM-PCA compares to the Mix-PCA (blue) or to the HMM-Gaussian (red). Therefore, the points that lie to the left of the diagonal line correspond to simulations where the HMM-PCA performed better than the other models, and the points that lie to the right represent simulations where the HMM-PCA performed worse. Given that *K* = 2, and because the order of the states is non-identifiable, this measure of accuracy ranges between 0.5 and 1.0. Models with accuracy of around 0.5 correspond to degenerate solutions, where one of the two states was obliterated by the inference. A summary of the average accuracy is shown in the text boxes. As expected, HMM-PCA has almost perfect accuracy for the p0 = 2 cases, and deteriorates to a moderate extent when *p*_0_ = 3. Most importantly, HMM-PCA also outperforms HMM-Gaussian in most simulations, confirming that approaching the problem in one single step leads to superior solutions when there are time-varying FC information in low-order PCs; i.e. when time-varying FC modulations are subtle, as it is the case in in most fMRI data sets. HMM-PCA also performs better than the Mix-PCA, highlighting the importance of accounting for the temporal structure of the data. (Note that in real data the temporal structure is stronger than in these simulations, so this difference will be even larger).

**Figure 3:**
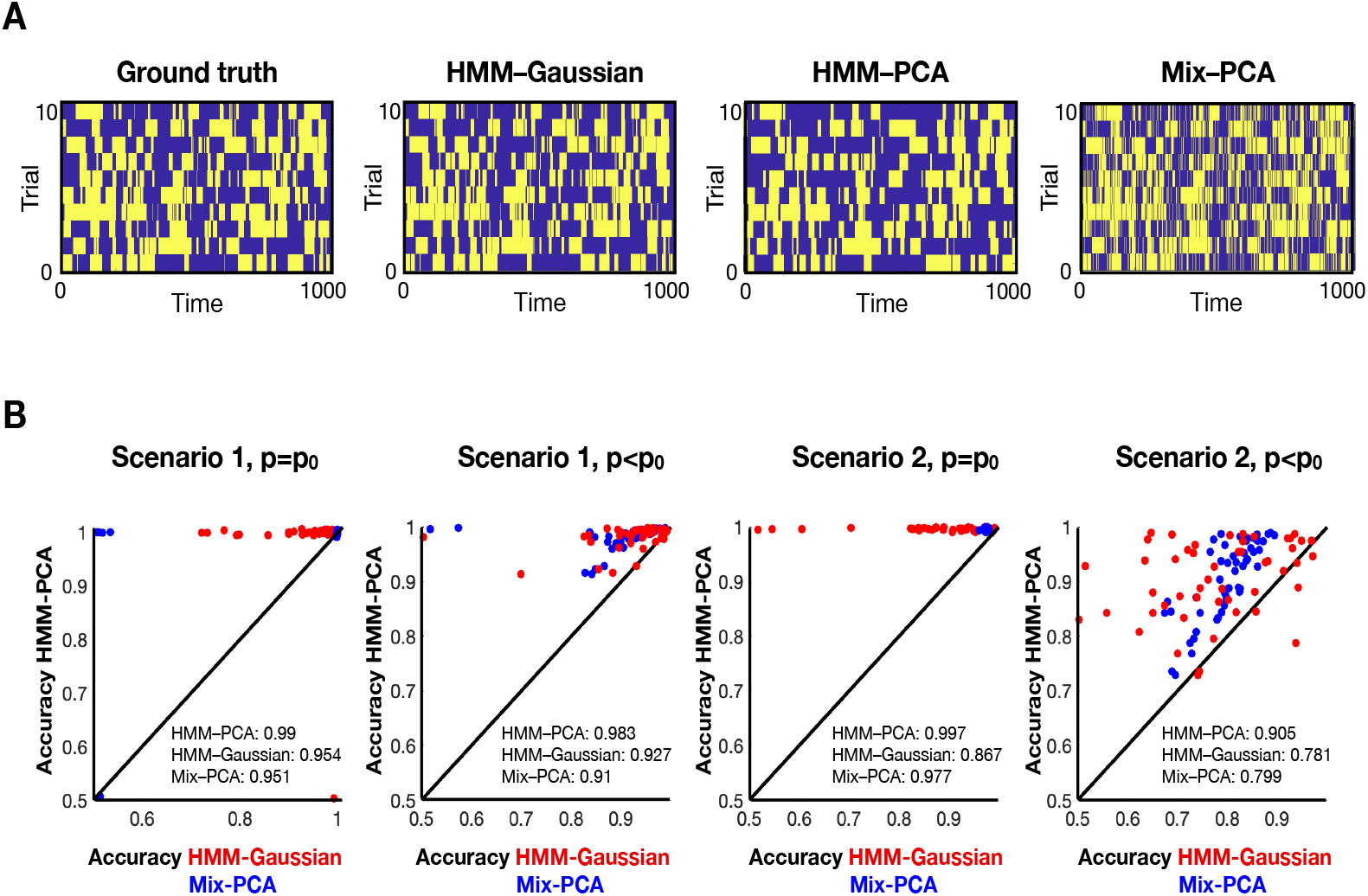
By performing the estimation in one single step and acknowledging the temporal nature of the data, HMM-PCA outperforms the HMM-Gaussian and Mix-PCA approaches **A**. Example of how the different models recover the ground-truth state time courses. **B**. Comparative accuracy between HMM-PCA (Y-axis), HMM-Gaussian (X-axis, red) and Mix-PCA (X-axis, blue). Each dot represents one repetition of the simulations, and accuracy is measured in terms of how well each method recovered the ground-truth state time courses.

## 3 Results on real fMRI data

So far, we have discussed the different models from a theoretical perspective and through simulations. Focusing on the HMM-based solutions, we now test the HMM-PCA and the HMM-Gaussian approaches on real resting-state fMRI data using 820 subjects from the HCP data set, where each subject underwent four 15-min sessions (TR=750ms) in the scanner. We used a data-driven parcellation obtained through spatial independent component analysis (ICA) with 50 components [3]. The time series of these ICA components were then standardised separately for each session, and then submitted to a stochastic variety of the HMM inference specially designed to deal with big volumes of data [14]. Since the estimation of the HMM parameters may return, for the same model and data, slightly different results for each run of the inference [24], we ran the inference five times per model, with *K* = 12 states each. In brief, we found that the main FC patterns were not anatomically very different between the two approaches, but that these differences were behaviourally informative, which can be taken as indirect evidence of the superior sensitivity of the HMM-PCA approach.

Asking whether the HMM-PCA states represent meaningful patterns of FC is not straightforward because there is no ground-truth available. Since both approaches are based on PCA, however, we expect that both approaches should be able to capture the main trends in the data to a relatively comparable extent. Given that HMM-Gaussian was shown to produce meaningful estimations in previous work [16, 8], proving that this is the case would situate HMM-PCA on first base. As an example, Figure 4A presents connectedness maps for two given states, where connectedness (or degree) is defined as how much each region correlates with the rest of the brain. The maps were centred across states, such that, if a region exhibits a positive value within a given state, then that region is more correlated to the rest brain’s voxels within this state than on average. One of the states is closely associated to the default mode network [2], and the other to the sensorimotor and visual systems. For these two states, both methods capture largely similar anatomical features. Based on the correlation between their FC patterns (specifically, by transforming the states’ covariance matrices into correlation matrices, taking the Fisher transformation, and then correlating the off-diagonal elements of these matrices between each pair of states), we then used the Hungarian algorithm [25] to match each HMM-PCA state to a HMM-Gaussian state. On the left, Figure 4B shows the resulting HMM-PCA vs HMM-Gaussian correlations for each pair of states, where the diagonal corresponds to states that were matched, and the off-diagonal to any other pair of states. On the right, Figure 4B shows the distribution of between-state correlations for states that were matched to each other (blue) and, to provide context, the correlations between states that were not matched (red). This result indicates that the main time-varying FC patterns are relatively preserved between the two methods.

**Figure 4:**
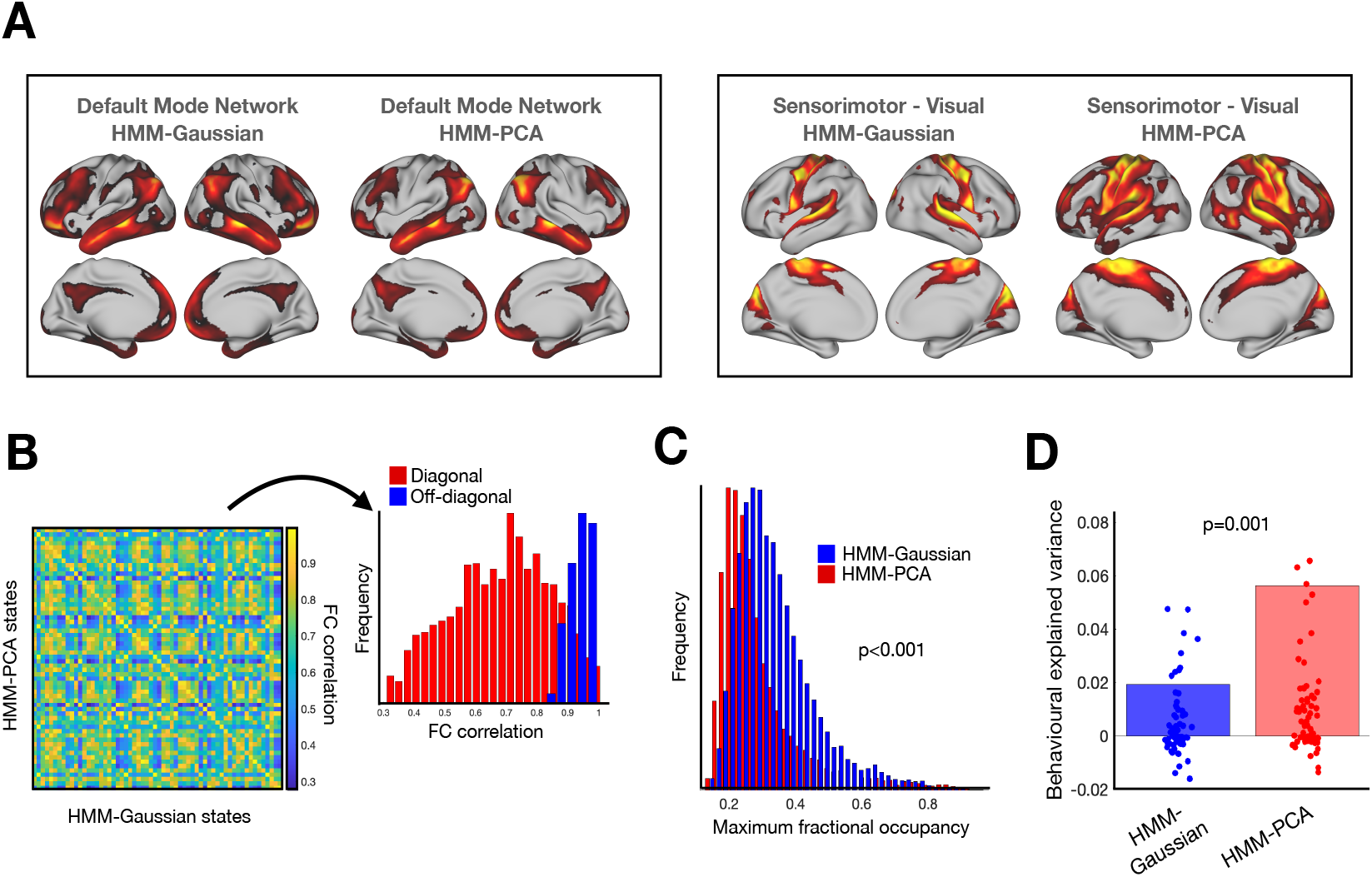
Comparison of HMM-Gaussian and HMM-PCA on resting-state data from 820 Human Connectome Project subjects [22]. **A**. Two example states per model, default mode network and sensorimotor-visual, where the maps reflect the degree, i.e. the total amount of connectivity between each voxel and the rest of the brain. **B**. The states are relatively comparable across the two models. Left: Correlation matrix between the HMM-PCA and the HMM-Gaussian states (in terms of Pearson’s correlation between the off-diagonal elements of the states’ FC matrices) across 5 runs of the inference. Right: Distribution of the diagonal elements (red) and off-diagonal (blue) elements, where the red elements reflect the similarity between corresponding states (i.e. an HMM-PCA state and an HMM-Gaussian state that correspond to each other), and the blue elements reflect the similarity between any two other states. **C**. HMM-PCA is more sensitive to time-varying FC than HMM-Gaussian, as reflected by the distribution of maximum fractional occupancies: HMM-PCA has a more distributed representation of the sessions in terms of the used states, while for the HMM-Gaussian solutions it is more frequent that one single state occupies most of the session. **D**. HMM-PCA produces solutions that are more explanatory of behaviour. Using the fractional occupancy across runs (i.e. the amount of time spent on each state), the HMM-PCA models are better able to predict behavioural traits across a range of demographical, intelligence, personality and affective-related variables; each dot represent the explained variance for the cross-validated prediction of one behavioural trait, and the bars represent the average across traits.

In light of this result, we sought to investigate whether the theoretical benefits of HMM-PCA have a practical impact on real data. First, for each run of the inference, we extracted the fractional occupancies, defined as the percentage of time spent on each state for each session; then, we computed the maximum fractional occupancy, a value assigned to each session reflecting the percentage of occupancy of the state that is most active within that session (for example, a maximum fractional occupancy of 1.0 means that one single state is active during the entire session). Using these, Figure 4C shows that HMM-PCA produced solutions with a higher sensitivity to time-varying FC, where it was less frequent for one single state to dominate an entire session and sessions were more distributed in terms of their state representation (p-value<0.001, permutation testing). We then used the 12 × 5 fractional occupancy values as features to predict a collection of behavioural traits. In particular, we selected 63 traits across different domains including demographical, affective, personality- and intelligence-related [5], and performed cross-validated predictions respecting the family structure of the HCP data [26]. As shown in Figure 4D, the cross-validated predictions were found to be significantly more accurate for HMM-PCA than for HMM-Gaussian (p-value=0.001, permutation testing). Albeit in an indirect way, these results suggests that HMM-PCA might be better able to describe time-varying FC in high-dimensional fMRI data, and that the theoretical limitations discussed above might have an impact in practice.

## 4 Discussion

In this paper, we have addressed the question of how to estimate patterns of time-varying FC in high-dimensional fMRI data. We have shown that the standard approach of sequentially applying PCA and then feeding the resulting PCs to an HMM or Bayesian mixture model, although useful in practice, may suffer from biases and loss of sensitivity. On these grounds, we have introduced a new variety of the HMM, the HMM-PCA, where each state is a Bayesian PCA decomposition. Critically, the HMM states not only express a (linearly optimal) dimensionality-reduction of the data, but also encode a correlation pattern between regions. Therefore, by fusing dimensionality reduction and time-varying FC in one single step, this approach can bypass some of the discussed limitations. This was shown through simulations and also in real fMRI data, where the HMM-PCA states were significantly better able to predict a number of behavioural traits.

Note that, since PCA is computed on the full data covariance, it is theoretically possible for the states transitions to be driven by changes in the pattern of relative variance across channels with little or no contribution of the between-regions covariance. If this were true, however, we would see no meaningful patterns after normalising by the variance (i.e. after transforming the covariance matrices into correlation matrices). The connectivity maps shown in Figure 4A, as well as the between-state FC correlations shown in Figure 4B, demonstrate that this it not the case, and that there are within-session FC modulations driving the estimation. This is in agreement with our previous work on the HCP data, where we showed (i) that HMM states have unique information in the off-diagonal elements of the covariance matrix, and (ii) that HMM models based solely on the variance (i.e. with no covariance) did not predict behaviour as accurately as those that model the full covariance [5, 8].

Another important question is the role played by between-subject FC differences on the inference, given that the models are run at the group level with no explicit consideration of these differences. Sometimes, too large between-subject differences can overshadow within-session FC modulations, resulting in entire sessions being occupied by one single state. When this happens, the HMM does no longer capture time-varying FC. Here, we showed that, on HCP data at least, the HMM-PCA approach is better able to capture time-varying FC. More generally, the HMM can be used as a tool to diagnose whether time-varying FC is negligible or dominated by between-subject FC differences. If the latter turns out to be the case, the issue should probably be dealt with at the level of data preprocessing, for example improving the co-registration of the data [27].

Finally, a possibility not considered in this paper is to run the HMM-PCA approach directly on the original voxel space. Because of the very large dimensionality of the volumetric and surface spaces in fMRI, this approach would be too computationally demanding and additional mechanisms would be needed to avoid overfitting due to the large number of parameters in each state.

## Acknowledgments and Disclosure of Funding

I thank Angus Stevner, Christine Ahrens, Piergiorgio Salvan, Mark Woolrich and Steve Smith for conversations that motivated this work. The study was supported by a Novonordisk Foundation grant NNF19OC-0054895. I declare no competing interests.

